# RNA editing enzyme APOBEC3A promotes pro-inflammatory (M1) macrophage polarization

**DOI:** 10.1101/2020.03.20.988980

**Authors:** Emad Y. Alqassim, Shraddha Sharma, Anm Nazmul H. Khan, Tiffany Emmons, Eduardo Cortes-Gomez, Kelly L. Singel, Jaron Mark, Bruce A. Davidson, A. J. Robert McGray, Qian Liu, Brian D. Lichty, Kirsten B. Moysich, Jianmin Wang, Kunle Odunsi, Brahm H. Segal, Bora E. Baysal

## Abstract

Pro-inflammatory (M1) macrophage polarization is associated with microbicidal and antitumor responses. We recently described APOBEC3A-mediated cytosine-to-uracil (C>U) RNA editing during M1 polarization. However, the functional significance of this editing is unknown. Here, we find that APOBEC3A-mediated cellular RNA editing can also be induced by influenza or Maraba virus infections of normal human macrophages, and by interferons in tumor-associated macrophages. Gene knockdown and RNA_Seq analyses show that APOBEC3A mediates C>U RNA editing of 209 exonic/UTR sites in 203 genes during M1 polarization. The highest level of deleterious C>U RNA editing occurred in *THOC5*, encoding a nuclear mRNA export protein implicated in M-CSF-driven macrophage differentiation. Knockdown of APOBEC3A reduces pro-inflammatory M1 markers including *IL6, IL23A* and *IL12B* gene expression, CD86 surface protein expression, and TNF-α, IL-1β and IL-6 cytokine secretion, and increases glycolysis and glycolytic capacity. Thus, APOBEC3A cytidine deaminase plays an important role in transcriptomic and functional polarization of M1 macrophages.

## Introduction

Macrophages are tissue-localized myeloid cells that originate during embryonic development or from recruited circulating monocytes. Monocytes mature into macrophages, which have specialized functions depending on the tissue they are settled. Macrophages are known to be active secretory cells that regulate host defense, inflammation, and homeostasis. They are also antigen presenting cells which are involved in initiating specific T cell responses (Laskin 2009). Through phagocytosis, macrophages contain bacterial and fungal infection, clear debris and apoptotic cells, and modulate anti-viral and anti-tumor immunity (Zhang and Mosser 2008). Macrophages have a wide range of diversity in their phenotype and functions in response to environmental cues, such as microbial products, products of cellular injury, cytokines, and hypoxia (Wynn et al., 2013). The biological functions of macrophages are mediated by specific subpopulations that are polarized phenotypically by exposure to specific mediators. Interferon-γ (IFN-γ), TNF-α, or pathogen-associated molecular patterns (PAMPs) activate pro-inflammatory M1 or classical polarization. M1 macrophages have potent microbicidal activity and release pro-inflammatory cytokines such as IL12 and TNF-α. (Edwards et al., 2006; Van Ginderachter et al., 2006). M1 macrophages augment T cell immunity and control intracellular infections (e.g., mycobacteria and Listeria), and enhance anti-tumor immunity. Alternatively activated (M2) macrophages are involved in anti-inflammatory and immunosuppressive activity through the release of IL10 and synthesis of mediators (e.g., VEGF and Arginase-1) that promote angiogenesis, tissue remolding and wound repair. M2 macrophages can be induced by fungal cells, various parasites, allergy, interleukin-4 (IL4), IL10, IL13 and tumor growth factor beta (TGF-B) (Porcheray et al., 2005; Röszer 2015; Krishnan et al., 2018). While macrophages are more likely to exist on a broad spectrum between the M1/M2 definitions, the M1 classification can be used to broadly describe inflammatory cells. Macrophage polarization and plasticity are important in normal physiology and in disease pathogenesis, and are exploitable therapeutically, for example, enhancing M1 polarization to augment intracellular antimicrobial defense and antitumor immunity.

The phenotype switch of macrophage is still not fully understood, but evidence points to mitochondrial function, reactive oxygen species, and metabolic changes regulating this switch. In general, M1 macrophages depend on glycolysis for ATP production, produce more reactive oxygen species (ROS) and accumulate succinate compared to resting macrophage (Mills et al., 2016; O’Neill and Pearce 2016). M1 polarization is generally characterized by induction of aerobic glycolysis along with high glucose uptake and high pyruvate-to-lactate conversion rate. On the other hand, M2 macrophages use oxidative phosphorylation as a permanent source of energy with a lower rate of glycolysis compare to M1 macrophages. Furthermore, forcing oxidative phosphorylation in M1 polarized macrophages can drive them to be M2 macrophages (Vats et al., 2006; Rodríguez-Prados et al., 2010; Galván-Peña and O’Neill 2014). Succinate dehydrogenase is well known to be a critical regulator of inflammatory macrophages. Following LPS stimulation, macrophages shift to aerobic glycolysis (Warburg effect) and production of pro-inflammatory cytokines (e.g., IL-1ß). LPS stimulation activates mitochondrial succinate dehydrogenase (SDH) resulting in increased succinate oxidation and ROS production at the expense of ATP synthesis, while inhibition of succinate oxidation dampened inflammation and was protective in LPS shock. (Mills et al., 2016). These results highlight the importance of SDH in metabolic programming of macrophages and the need to understand mechanisms that regulate its expression and function.

Macrophage responses to environmental cues (e.g., LPS, hypoxia, cytokines) are modulated by complex signaling pathways that include transcriptional regulation (Lawrence and Natoli 2011). microRNA (miRNAs), DNA methylation and histone modification have all been reported to influence macrophage polarization (Zhou et al., 2017). In addition to these well-characterized pathways for epigenetic modification, enzyme-regulated RNA editing may be another mechanism regulating macrophage responses. RNA editing is a posttranscriptional mechanism that alters transcript sequences, without any change in the encoding DNA sequences and therefore can generate protein diversity (Farajollahi and Maas 2010). By changing RNA sequences, RNA editing can generate protein diversity in single-cell organisms, plants, and animals (Farajollahi and Maas 2010). Two significant types of RNA editing have been recognized in mammals. One type involves adenine conversion to inosine (A>I) by adenosine deaminase enzymes and occurs in hundreds of thousands of sites located mostly in non-coding, intronic regions (Eisenberg and Levanon 2018). The other dominant type of RNA editing is C>U RNA editing which is catalyzed by the apolipoprotein B-editing catalytic polypeptide-like (APOBEC) family of cytidine deaminase enzymes (Bransteitter et al., 2009; Salter et al., 2016).

The initially described activity of APOBEC-mediated C>U editing involved site-specific mRNA editing of ApoB, a protein required for the assembly of very low-density lipoproteins from lipids by APOBEC1 (Teng et al., 1993). APOBEC3 proteins have a well-recognized role in inhibiting retroviruses, endogenous retroelements and other viruses in *in vitro* systems. Such inhibitions may be dependent or independent of the deaminase functions of APOBEC3s (Holmes et al., 2007). Recently, we identified the cellular RNA editing functions of APOBEC3A (A3A) (Sharma et al., 2015) and APOBEC3G (A3G) (Sharma et al., 2019). A3A and A3G each target specific cytidines located in stem-loop structures in distinct set of transcripts (Sharma and Baysal 2017). RNA editing by A3A and A3G can be induced by exogenous over-expression in cell lines (Sharma et al., 2016; Sharma et al., 2017a; Grünewald et al., 2019). A3A-mediated RNA editing is endogeneously induced by IFN-γ during M1 macrophage polarization and by type 1 interferons (IFN-1) in monocytes/macrophages (Sharma et al., 2015). In addition to IFNs, hypoxia and cellular crowding induces endogenous C>U RNA editing by A3A in monocytes and by A3G in natural killer (NK) cells (Baysal et al., 2013; Sharma et al., 2015; Sharma et al., 2019). Hypoxia and IFN-1 additively enhance A3A-mediated RNA editing in monocytes, leading to RNA editing levels of ~ 80% in transcripts of certain genes. A3A-or A3G-induced RNA editing by cellular crowding and hypoxia can be mimicked by the inhibition of mitochondrial respiration (Sharma et al., 2017b; Sharma et al., 2019). A3G-mediated RNA editing triggers a Warburg-like metabolic remodeling in HuT78 T cell lymphoma line (Sharma et al., 2019). These findings suggest that APOBEC3-mediated RNA editing may play a role in hypoxia-or IFN-induced cell stress response.

Humans have ten APOBEC genes, but only A3A mediates RNA editing in monocytes and macrophages in scores of genes’ transcripts, including the c.136C>U (R46X) event in the succinate dehydrogenase B *(SDHB)* mRNA (Baysal 2007). In this study, we aimed to delineate the role of A3A on macrophage function during M1 polarization and viral infection. We hypothesized that A3A plays an essential role in macrophage functions during M1 polarization and in response to viral infections. Using primary human monocyte-derived macrophages, we observed that A3A-mediated RNA editing during viral infections and M1 polarization targeted a broad range of genes, and had broad effects on transcriptome, pro-inflammatory and metabolic responses that drive M1 polarization. These results set the foundation for investigating the role of A3A in patients during sepsis and other diseases associated with pathologic macrophage responses.

## Results

### A3A induces *SDHB* RNA editing in M1 macrophages

We previously showed that IFN-γ and LPS induce RNA editing in M1 macrophages and that the level of editing in several sites is reduced with KD of A3A (Sharma et al., 2015). Here, M0 cells were treated for two days with 20 ng/ml recombinant human IFN-γ (Life Technologies) and 50 ng/ml Escherichia coli LPS, or 20 ng/ml recombinant human IL4, for M1 or M2 polarization, respectively. For knockdown (KD) experiments, a day before induction of M1 polarization, M0 macrophages were transfected with negative control scrambled (SC) or A3A siRNAs. We first confirmed that the induction of *SDHB* c.136C>U RNA editing occurs in M1 macrophages but not in M0 or M2 macrophages **(Figure 1A)**. To further confirm whether A3A mediates *SDHB* C.136C>U RNA editing in M1 macrophages, we knocked down A3A by siRNA and induced the polarization of M0 macrophages to M1 macrophages with LPS and IFN-γ. Knockdown of A3A led to a significant reduction in the level of *SDHB* C.136C>U RNA editing as tested in 8 additional donor samples **(Figure 1B)**. These findings demonstrate that A3A is responsible for the observed *SDHB* c.136C>U RNA editing in M1 macrophages. Macrophage viability, assessed by flow cytometry, was similar in A3A KD and A3A SC M1 macrophages **(Supp. Figure 1).**

**Figure 1.**
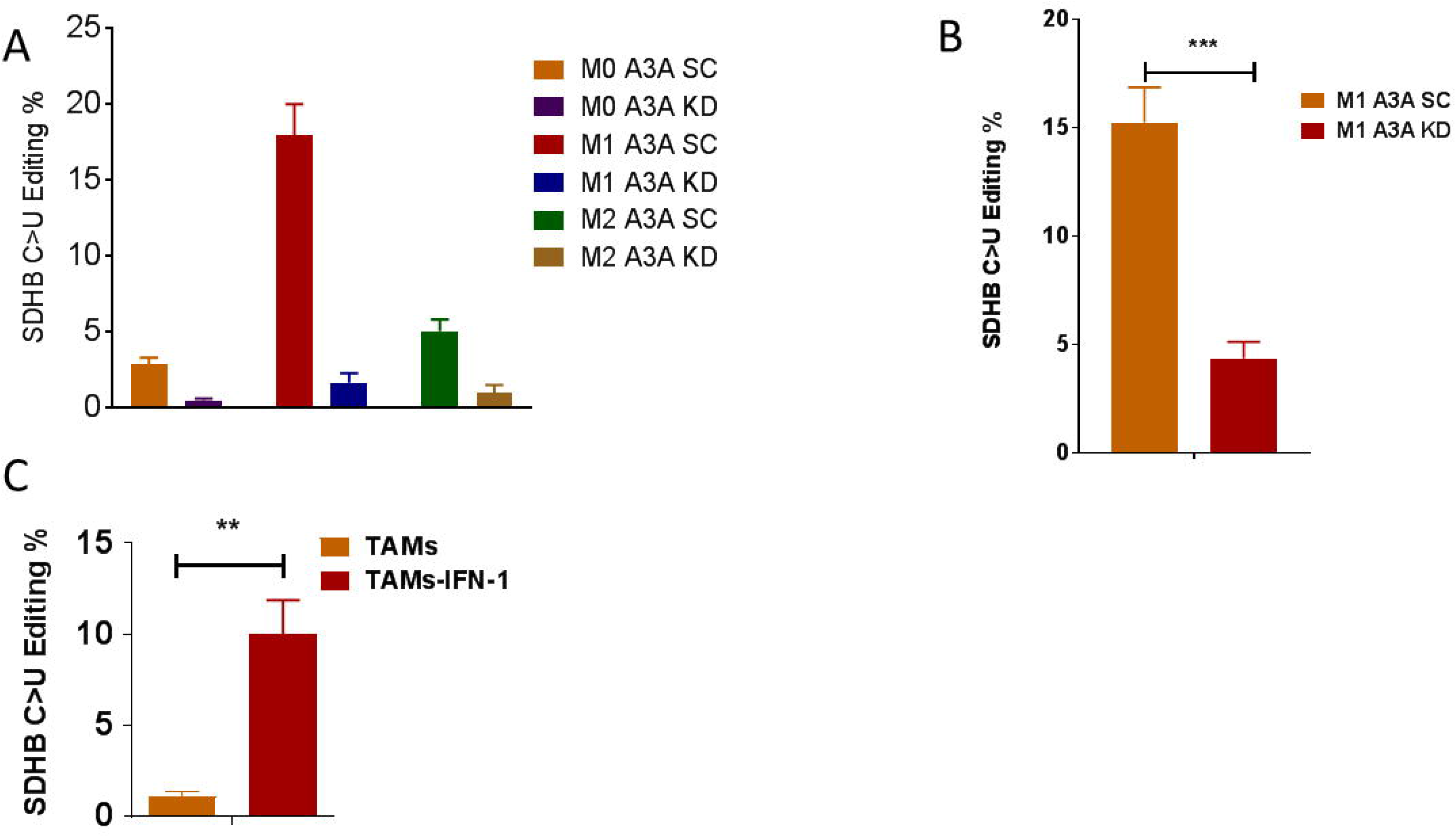
*SDHB* c.C136U RNA editing is induced in M1 macrophages and mediated by A3A. **A)** RNA editing was induced in M1 macrophages but not in M0 or M2 macrophages (n=2 replicates from 1 representative donor). **B)** A3A was required for *SDHB* RNA editing. P = 0.0001, Paired t test, two-tailed, n=8 donors. **C)** IFN-1 (IFN-α, 1,000 U/ml) induces RNA editing in ovarian tumor associated macrophages (TAMs) from ascites of ovarian cancer patients. P = 0.0084, Paired t test, two-tailed, n=5 donors. Data shown are mean ± SEM.

### IFN-1 induces RNA editing in tumor-associated macrophages (TAMs)

After showing the induction of A3A-induced RNA editing in normal macrophages in the context of M1 polarization, we studied TAMs. Although TAMs are M2-like, the tumor microenvironment is often hypoxic and characterized by pro-inflammatory cytokines, including interferon. Hence, we asked whether TAMs can be repolarized to M1-like phenotype by analyzing A3A-mediated RNA editing. We investigated the impact of IFN-1 on TAMs purified from the ascites of 5 patients with newly diagnosed epithelial ovarian cancer. TAMs were isolated by CD14 microbeads, and purity was confirmed by CD33^hi^ expression and by cytology. Compared to untreated samples, we found that editing level was induced significantly in TAMs treated with IFN-1 (**Figure 1C**). These results show that A3A-mediated RNA editing is also inducible in TAMs by IFN-1.

### Viral infections induce RNA editing by A3A in primary human M0 macrophages

We next tested whether viral infection, which triggers IFN production in a more physiologically relevant setting, would induce A3A-mediated RNA editing in M0 macrophages. Normal donor monocyte-derived macrophages were generated as described above and infected with two available viruses: Maraba and influenza. Maraba is a genetically modified SS-RNA oncolytic rhabdovirus being developed for cancer therapy (Brun et al., 2010; Tong et al., 2017). Influenza is a SS-RNA orthomyxovirus. We observed that viral infection induced *SDHB* RNA editing **(Figure 2A)**. Then we tested whether induction of RNA editing in M0-infected was A3A-dependent. Normal donor monocyte-derived macrophages were generated as described above and A3A was knocked down 24h before infection with Maraba or influenza virus. We observed that RNA editing level was suppressed in M0 A3A KD compared to M0 A3A SC **(Figure 2A).** Using ELISA, a higher concentration of IFN-1 was detected in the supernatant of M0 macrophages infected with Maraba virus compared to non-infected M0 macrophages **(Figure 2B)**. These results show that viruses can induce A3A-mediated RNA editing in macrophages, likely mediated by increased IFN production that is known to induce A3A expression levels by hundreds-fold (Sharma et al., 2015).

**Figure 2.**
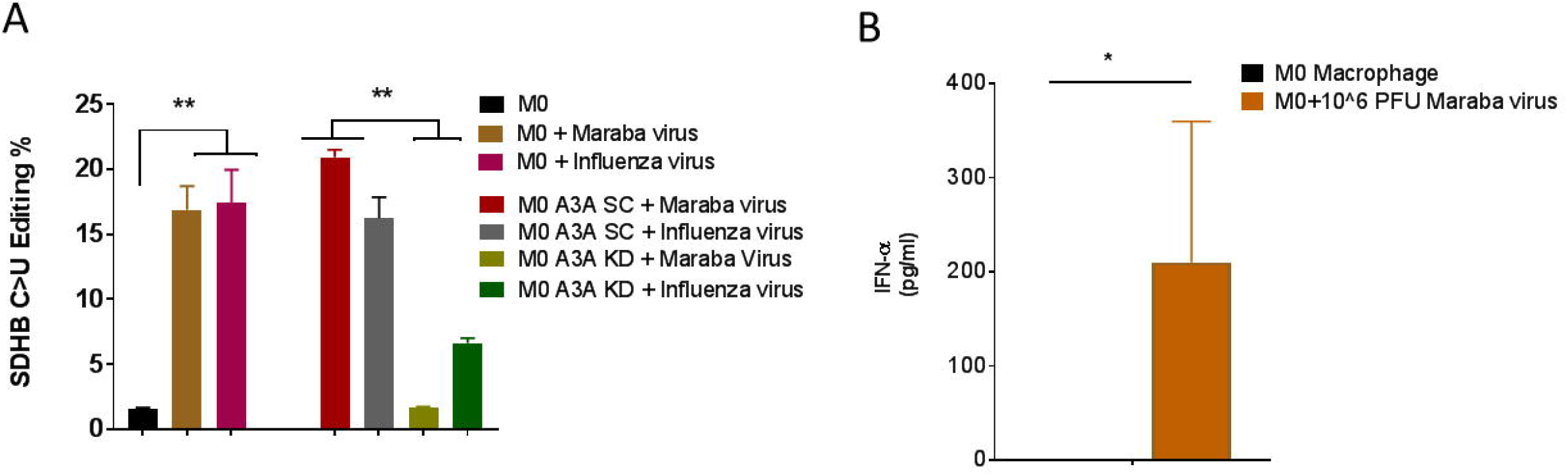
Viruses induce A3A-mediated RNA editing in macrophages. Circulating healthy donor monocytes (~1X10^6^ cells per well) were differentiated into M0 macrophages followed by A3A KD or scramble (SC). After 24h, were infected with Maraba (10^6 PFU per well) or influenza virus (2×10^4^ PFU per well, MOI=0.02), and *SDHB* RNA editing was assessed at 48h after infection: n=3 biological replicates for Maraba and n=2 biological replicates for influenza viruses. **A)** Viral infection of M0 macrophages increased *SDHB* RNA editing levels for both viruses. **p=0.0095, Mann-Whitney test, two-tailed. A3A KD significantly reduced *SDHB* RNA editing levels induced by A3A SC for both viruses. **p=0.0079, Mann-Whitney test, two-tailed. **B)** IFN-α levels in culture supernatant significantly increase after Maraba virus infection. *p=0.0286, Mann-Whitney test, two-tailed, n=3 donors. Data shown are mean ± SEM.

### APOBEC3A catalyzes the majority of C>U RNA editing events during M1 polarization

We generated monocyte-derived macrophages from 3 normal donors and induced M1 polarization in cells transfected with A3A KD and A3A control SC siRNAs. Non-polarized macrophages (M0) were used as specificity controls. RNA editing of *SDHB* c.C136C>U was first confirmed by RT-PCR and ranged between 11-21% in SC M1-versus 1-3% in A3A KD M1-polarized macrophages (RNA editing in M0 macrophages was 0-2%). Paired RNA_Seq analysis was performed comparing SC and A3A KD M1-polarized macrophages from the same donor.

To directly test the extent that A3A catalyzes these RNA editing events, we first determined the C>U RNA editing events that were induced at least 2-fold in M1 SC siRNA relative to the control M0 samples and that occur at >5% level in any experimental group, using high-coverage RNA_Seq data and stringent filtering steps described in Methods. This analysis revealed 209 C>U RNA editing sites in 203 genes in SC M1-macrophages (**Supp. Table 1**). KD of A3A during M1 polarization reduced RNA editing levels by more than 2-fold in 180 of 209 (~86%) sites, indicating that A3A catalyzes the majority of RNA editing sites during M1 polarization **(Figure 3A)**. About half of the C>U RNA editing events catalyzed by A3A were synonymous (98/180=54.4%), followed by non-synonymous (36/180=20%) and 3’-UTR (33/180=18.3%) **(Figure 3B and 3C**). The RNA editing sites that were not significantly affected by A3A KD may represent genomic SNVs, true RNA-edited sites that are not catalyzed by A3A, or false positives.

**Figure 3.**
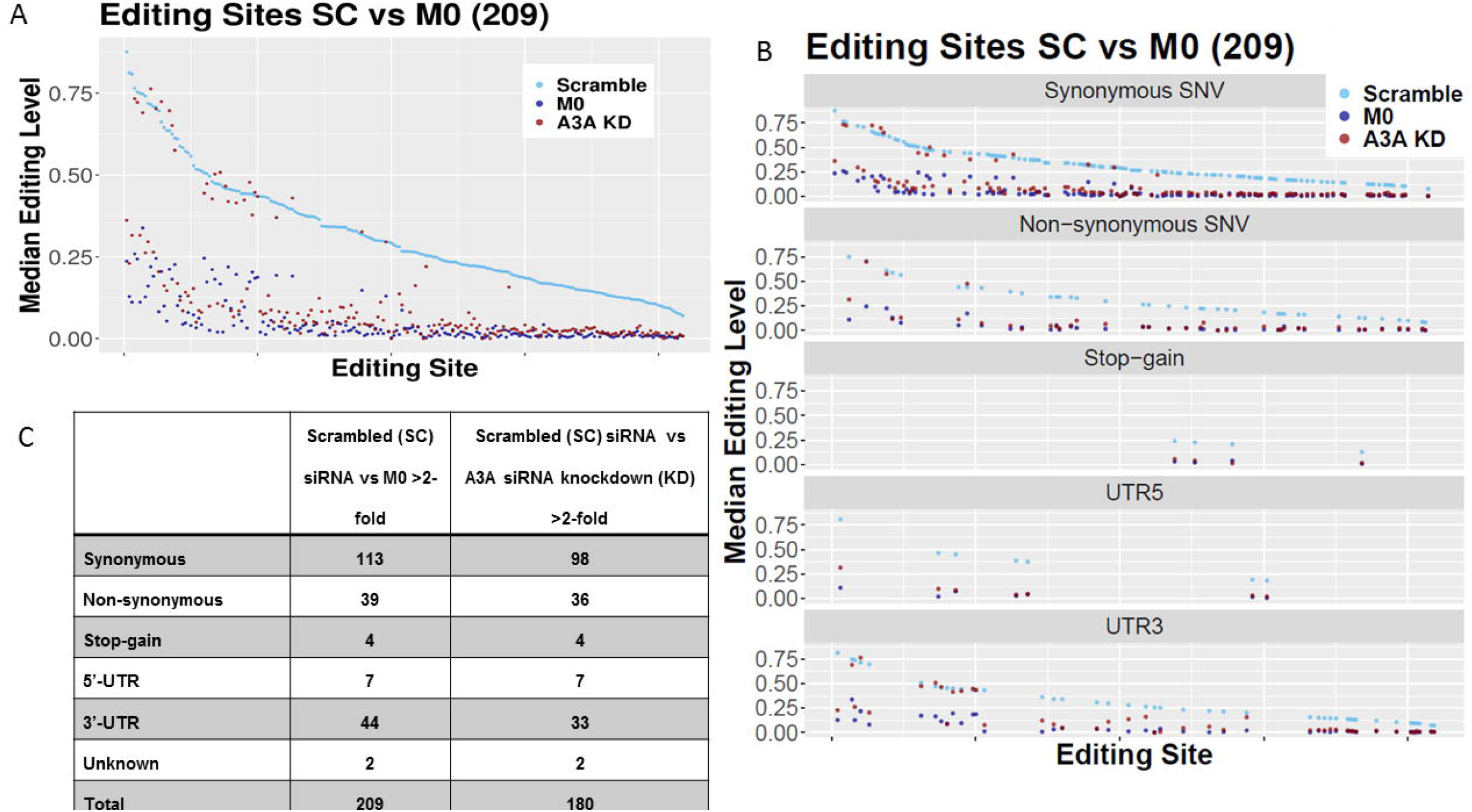
A3A catalyzes the majority of C>U RNA editing events during M1 polarization. **A)** A3A KD reduces the C>U RNA editing levels in most sites in M1 macrophages close to levels seen in M0 unpolarized macrophages. C>U RNA editing sites are ranked by their levels in M1 A3A SC (scramble) control. **B and C)** C>U RNA editing events in M1 A3A SC, M1 A3A KD and M0 cells are separately shown for variant types in graph.

Examination of the non-synonymous/stop-gain RNA editing events (**Supp. Table 1**) showed the highest editing level in *THOC5* with 75% RNA editing level in M1 A3A SC macrophages. The mean RNA editing level of *THOC5* was significantly reduced to about 32% in M1 A3A KD macrophages **(Table 1)**. *THOC5* RNA editing (c.C2047T:p.R683C) alters the last amino acid arginine, which is conserved from lamprey to human, to cysteine, and is predicted to be deleterious. Other genes including *ZNF124, ARSB, ATXN2, SLC37A2, GAA, PCGF3, CDYL2,* and *S1PR2* also showed high levels (>20%) of non-synonymous RNA editing in M1 A3A SC compared to M1 A3A KD macrophages **(Table 1).**

**Table 1:** High levels of non-synonymous C>U RNA editing by APOBEC3A during M1 polarization

| Gene | Genomic<br>coordinate of RNA<br>editing<br>(GRCh38/hg38) | RNA editing event | SC Average<br>Editing<br>Level | A3A KD<br>Average<br>Editing Level | SC/KD<br>Ratio |
| --- | --- | --- | --- | --- | --- |
| ***SDHB*** | chr1:17044825 | NM_003000:exon2:c.<br>C136T:p.R46X | 0.227 | 0.040 | 5.70 |
| ***THOC5*** | chr22:29508462 | NM_001002879:exon<br>20:c.C2047T:p.R683C | 0.750 | 0.316 | 2.38 |
| ***ZNF124*** | chr1:247157042 | NM_001297568:exon<br>4:c.C580T:p.R194C | 0.589 | 0.117 | 5.04 |
| ***ARSB*** | chr5:78984968 | NM_198709:exon2:c.<br>C281T:p.S94L | 0.434 | 0.074 | 5.86 |
| ***ATXN2*** | chr12:111516348 | NM_001310121:exon<br>10:c.C866T:p.S289L | 0.396 | 0.049 | 8.04 |
| ***SLC37A2*** | chr11:125080652 | NM_198277:exon7:c.<br>C566T:p.S189F | 0.342 | 0.049 | 7.00 |
| ***GAA*** | chr17:80107625 | NM_001079804:exon<br>4:c.C761T:p.S254L | 0.341 | 0.021 | 16.39 |
| ***PCGF3*** | chr4:743578 | NM_006315:exon7:c.<br>C367T;p.R123W | 0.300 | 0.066 | 4.55 |
| ***CDYL2*** | chr16:80684900 | NM_152342:exon2:c.<br>C254T;p.S85L | 0.248 | 0.018 | 13.77 |
| ***S1PR2*** | chr19:10224709 | NM_004230:exon2:c.<br>C197T;p.S66L | 0.220 | 0 | N/A |

Sanger sequencing of RT-PCR products obtained from a different monocyte donor confirmed high-level (~50%) of RNA editing, which is reduced by A3A KD, in *THOC5* and *ZNF124* gene transcripts during M1 polarization. As expected, M0 macrophages showed no evidence of RNA editing **(Figure 4)**. Together, these results demonstrate that mRNAs of certain genes are substantially mutated by A3A-mediated RNA editing in M1-polarized macrophages.

**Figure 4.**
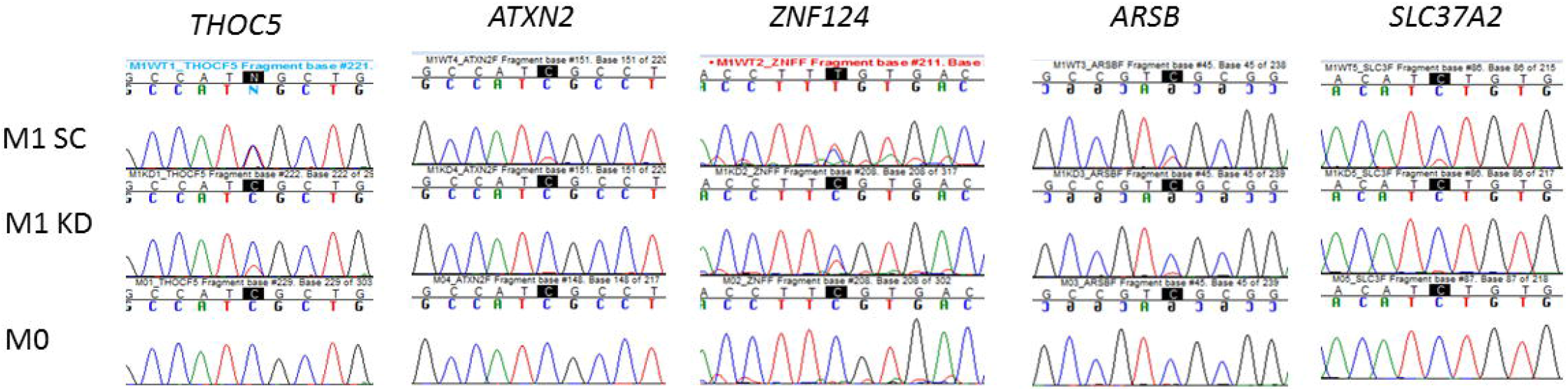
Sanger sequencing demonstrating A3A-mediated C>U RNA editing in select gene transcripts. Sanger sequencing of selected gene mRNAs confirms the induction of RNA editing in A3A M1 SC, which is reduced by A3A KD. Unpolarized M0 samples show no evidence RNA editing. The edited Cs are highlighted within black boxes.

### Effect of A3A KD on gene expression in M1 macrophages

Relative to A3A SC, A3A KD caused more than 2-fold downregulation of 81 genes and upregulation of 131 genes with statistical significance (Padj<0.05; **Supp. Table 2**). A3A was the second most downregulated gene upon A3A KD, confirming the intended siRNA targeting. Pro-inflammatory cytokines *IL6, IL23A* and *IL2B* were among the most downregulated genes upon A3A KD in M1 macrophages (**Figure 5A**). Examination of selected inflammation-related genes showed significant differences between M1 A3A SC and M1 A3A KD. A3A KD significantly reduced the expression of cell surface pro-inflammatory genes CD68, CD80 and CD86 but increased the expression of MRC1 (CD206), a marker of M2 polarization. The expression of inflammatory cytokine genes *TNF, IL6, IL23A, IL18* and *CXCL8* was also statistically significantly reduced by A3A KD **(Figure 5B)**. Gene ontology (GO) analysis showed that A3A augmented the expression of pathways driving cytokine/chemokine production and signaling, adaptive immunity, and cell death, but markedly down-regulated pathways involved in protein elongation/translation and metabolism and influenza viral RNA transcription and replication (Table 2). These results suggest that A3A plays an important role in the regulation of gene expression in M1 polarized macrophages, with the major effects being upregulation of proinflammatory genes and pathways while reducing genes for protein metabolism and influenza life cycle.

**Figure 5.**
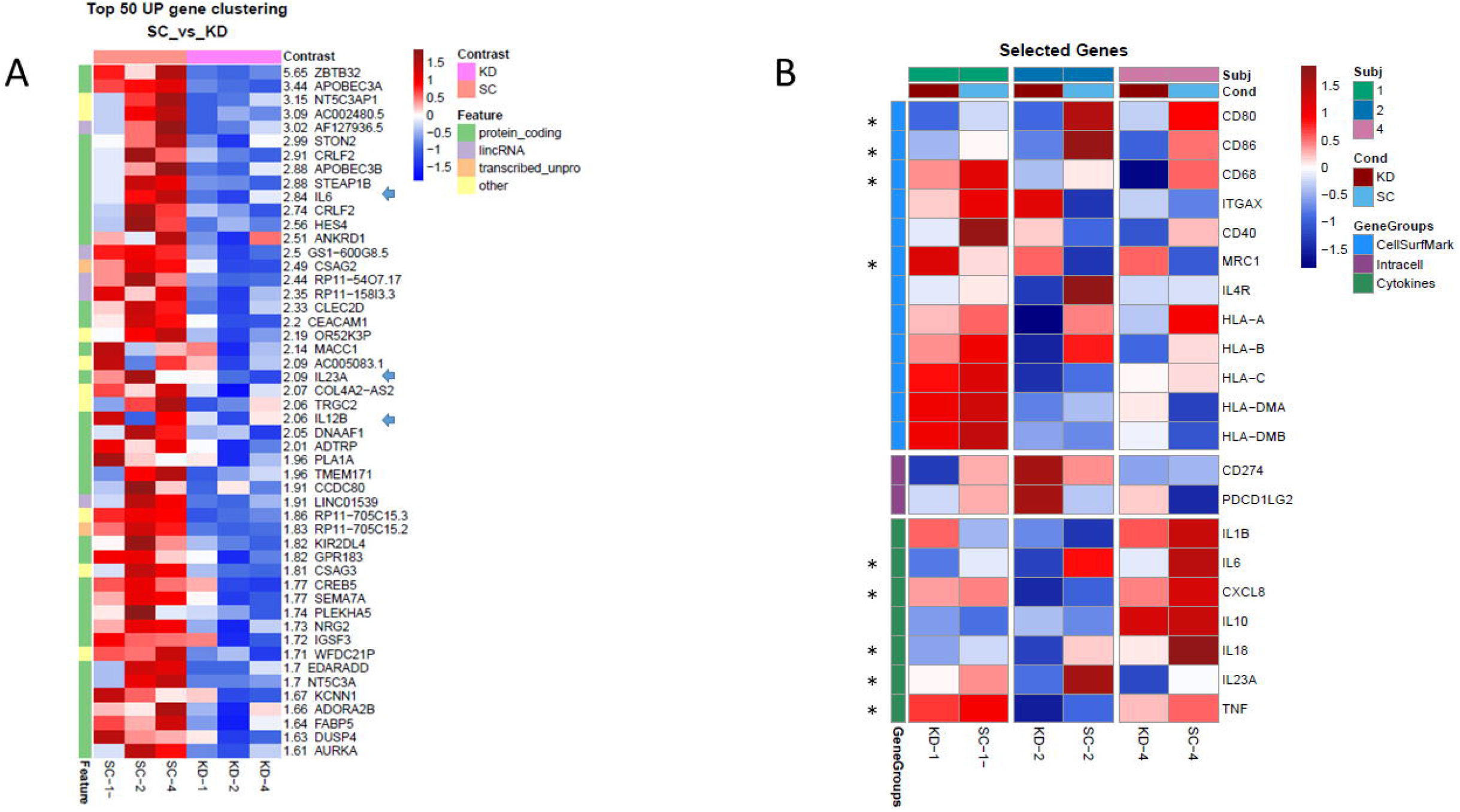
A3A augments inflammatory gene expression in M1-polarized macrophages. Heatmaps of selected pro-inflammatory genes show greater expression in M1 A3A SC macrophages compared to M1 A3A KD macrophages. **(A)** Heatmap of showing the top 50 most upregulated genes in M1 SC samples relative for M1 A3A KD samples include pro-inflammatory genes *IL6, IL23A* and *IL12B* (arrowheads). **(B)** Expression of *CD80, CD86, CD68, TNF, IL23A, IL18, CXCL8* and *IL6* genes are reduced with A3A KD (* p = <0.05), while MRC1 (CD206), which is associated with M2 polarization, had increased expression with A3A KD.

**Table 2:** Pathways Upregulated and Downregulated by A3A in M1-polarized macrophages.

| **Upregulated** | **Downregulated** |
| --- | --- |
| Adaptive immune system **** | Peptide elongation/translation **** |
| Cytokine-cytokine receptor **** | Glycolysis-gluconeogenesis * |
| G-protein coupled receptor **** | Pyruvate metabolism * |
| Interferon alpha-beta **** | PID-HIF1-TF pathway * |
| Antigen processing and presentation **** | Mitochondrial protein ** |
| Class-1- MHC mediated **** | Protein metabolism **** |
| Apoptosis ** | Pentose Phosphate Pathway * |
| Chemokines receptors **** | Arginine and Proline metabolism * |
| Stress pathway * | Influenza life cycle **** |
| Cytokines pathway ** | Influenza viral RNA transcription and replication **** |
| MHC- Class-II antigen presentation*** | Integrin-1 pathway ** |
| Death pathway * | Transport of glucose* |

FDR q-val by GO analysis comparing A3A SC versus A3A KD: \* <0.05; \*\* <0.01; \*\*\* <0.001; \*\*\*\* <0.001

### A3A enhances the release of inflammatory cytokines and CD86 expression in M1 macrophages

To examine whether the pro-inflammatory role of A3A observed at the gene expression level also extends into protein expression level, we evaluated the role of A3A in macrophage surface antigen expression and cytokine responses. The knockdown of A3A is confirmed by reduction in *SDHB* c.C136C>U RNA editing levels (**Figure 1**). Cytokine production by M1 A3A SC and M1 A3A KD macrophages was assessed by ELISA. A3A knockdown decreased TNF-α, IL-1ß, and IL6 secretion by M1 macrophages (**Figure 6A-C**).

**Figure 6.**
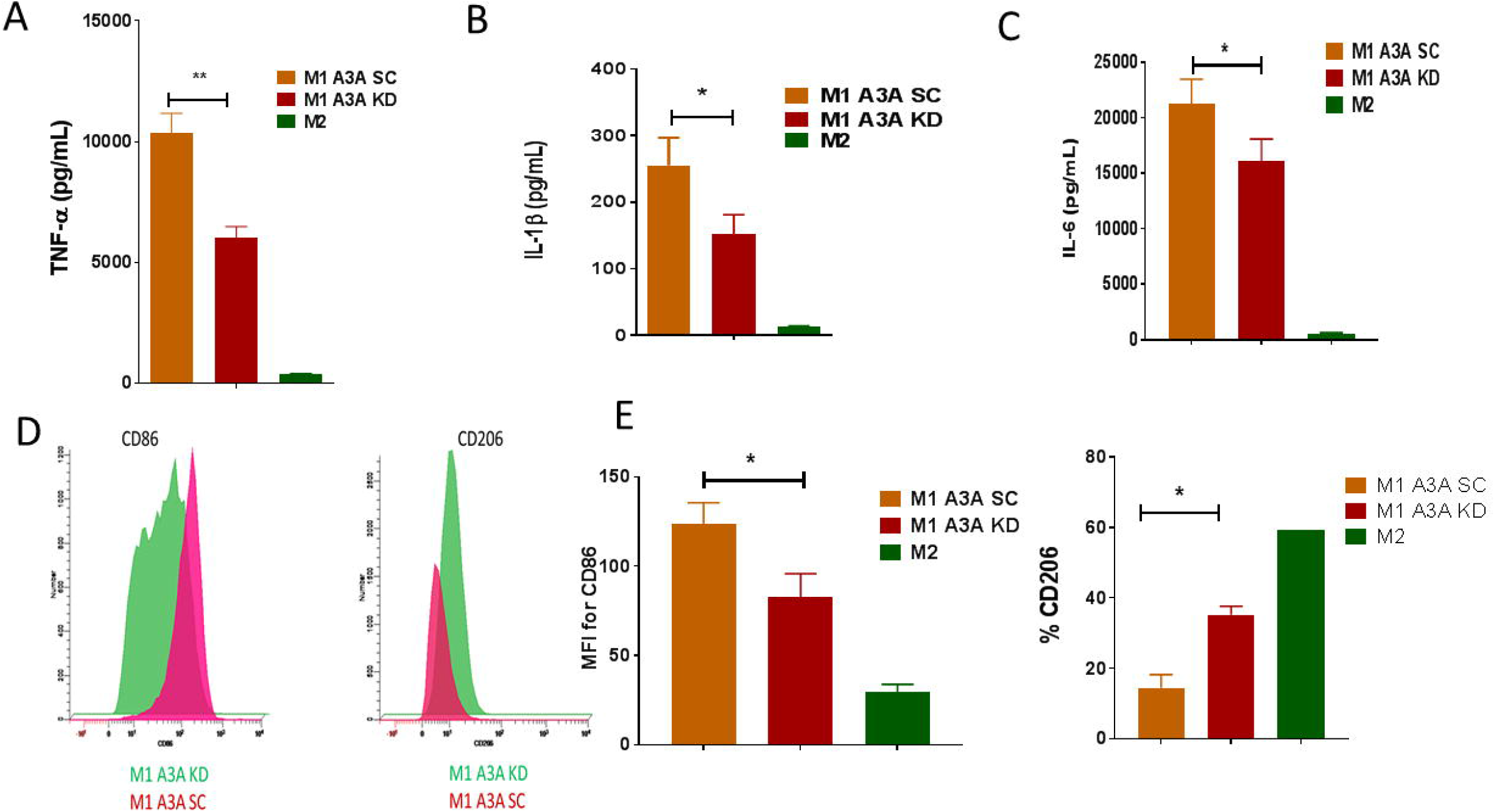
Knockdown (KD) of A3A impairs pro-inflammatory phenotype of M1 macrophages. **A)** A3A KD decreases TNF-α secretion by M1 macrophages from normal donors compared to controls (SC). P=0.0049, Paired t test, two-tailed, n= 5 donors **B)** A3A KD decreases IL-1ß secretion by M1 macrophages from normal donors compared to controls (SC). P=0.0103, Paired t test, two-tailed, n=6 donors **C)** A3A KD decreases IL6 secretion by M1 macrophages from normal donors compared to controls (SC). P=0.0192, Paired t test, two-tailed, n= 6 donors **D)** A3A KD reduces CD86, but enhances CD206 (MRC1) surface protein expression in M1-macrophages by flow cytometry. **E)** The quantification of CD86 (P=0.0313, Wilcoxon matched-pairs signed rank test, two-tailed, 6 donors) and CD206 (representative results from 2 donors shown) expression is made following M1 polarization in CD33-positive cells. MFI=Mean Fluorescent Intensity. Data shown are mean ± SEM.

We next next assessed by flow cytometry the role of A3A in modulating the surface expression of CD86 and CD206 as markers for M1 macrophages and M2 macrophages, respectively (Wolf et al., 2014). Compared to A3A SC macrophages, A3A KD M1 macrophages showed a reduction in CD86 expression (**Figure 6D, E**). CD206 expression was increased in M1 A3A KD compared to M1 A3A SC macrophages **(Figure 6D, E).** As expected, M2-polarized macrophages showed reduced CD86 and increased CD206 expression **(Figure 6E)**. These results demonstrated that A3A amplifies pro-inflammatory phenotype in M1 macrophages.

### A3A suppresses the level of glycolysis in M1 macrophages

While resting macrophages rely principally on mitochondrial respiration for energy, the switch to glycolysis under aerobic conditions (Warburg effect) is associated with polarization to M1 macrophages (Wolf et al., 2014). Since A3A plays a role in the augmentation of inflammatory responses of M1 macrophages, we hypothesized that A3A induces glycolysis during M1 polarization. To explore the impact of A3A on the metabolic phenotype of M1 macrophages, we evaluated glycolysis and mitochondrial respiration in A3A SC and A3A KD M1 macrophages by the SeaHorse Glycolytic Stress and Mitochondrial Stress assays, respectively. As expected, M1 macrophages had a higher glycolysis rate compared to M2 macrophages. Unexpectedly, the level of glycolysis was increased in A3A KD M1 macrophages compared to M1 A3A SC, indicating that A3A reduced glycolysis in M1 macrophages (**Figure 7A)**. These results are consistent with our GO analysis (Table 1) showing that A3A reduces expression of genes involved in glycolysis and other metabolic pathways. Basal oxygen consumption rates were similar between A3A KD M1 and A3A SC M1 macrophages (**Figure 7B**). These results suggest that A3A enhances the production of pro-inflammatory cytokines independently of the Warburg effect in M1 macrophages.

**Figure 7.**
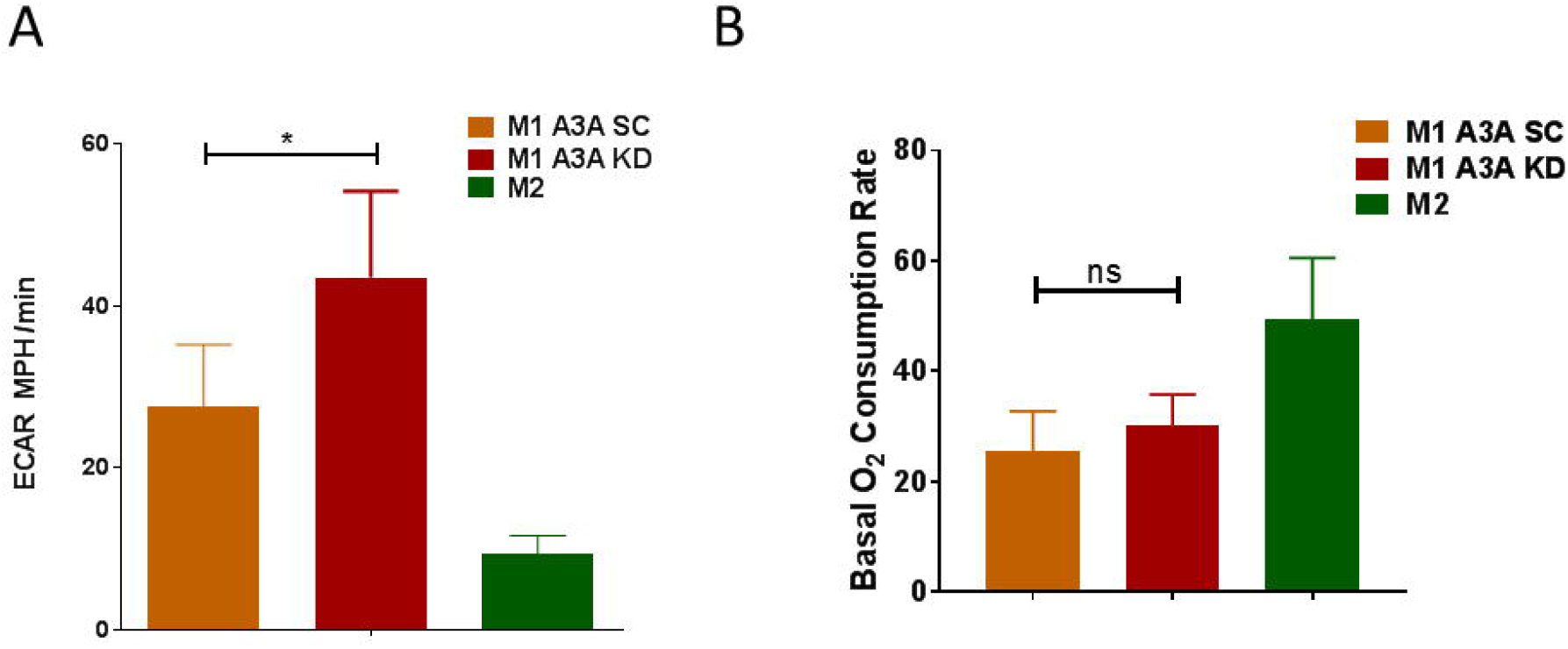
The role of A3A in glycolysis and oxygen consumption in macrophages. **A)** A3A KD enhances glycolysis (ECAR) compared to A3A SC in M1 macrophages derived from normal donor blood. P=0.0171, Paired t test, two-tailed, 5 donors **(B)** Basal oxygen consumption rate is similar between A3A KD M1 macrophages and A3A SC M1 macrophages. P not significant (P=0.686), Mann-Whitney test, two-tailed, 4 donors. Data shown are mean ± SEM.

## Discussion

Here, we provide evidence that A3A is required for transcriptomic and functional polarization of M1 macrophages, most likely through C>U RNA editing of scores of genes. We show that sitespecific C>U RNA editing by A3A can be induced by M1 polarization or viral infections in normal monocyte-derived macrophages, and by IFN-1 exposure in tumor-associated macrophages isolated from ovarian cancer-related ascites fluid. Together, these results point to a broad role for A3A in modulating macrophage responses to diverse conditions.

For the first time, to our knowledge, we observed that viruses also induce A3A-dependent cellular RNA editing, which in turn downregulates cellular genes for viral (influenza) replication (Table 2). Viral infections can drive M1 polarization that in turn enhances antiviral immunity. IFN-1 plays a role in M1 polarization of macrophages exposed to virus infection (Sang et al., 2015). Accordingly, IFN-1 concentration was detectable in the media of macrophages infected by the Maraba virus (**Figure 2B**). MG1 Maraba is an oncolytic virus (OV) that is being tested in phase1/2 clinical trials, and acts through multiple antitumor mechanisms including direct tumor cell lysis, modulation of tumor microenvironment to a more pro-inflammatory state and generation of tumor-specific T-cells (Martin and Bell 2018; Breitbach 2020). Our findings raise the speculation that when tumor-associated macrophages are infected by Maraba virus, A3A promotes their pro-inflammatory polarization, creating a favorable microenvironment for development of antitumor immunity, while at the same time enhancing antiviral response. Therefore, the role of A3A-mediated RNA editing in OV cancer therapy requires further studies.

A3A amplifies M1 macrophage polarization and pro-inflammatory cytokine responses. RNA-Seq analysis shows that A3A increased gene expression of pro-inflammatory cytokines like *TNF, IL6, IL18,* and *CXCL8,* while also limiting the glycolysis in M1 macrophages. GO analysis shows that A3A down-regulates glycolysis pathway in addition to all metabolic pathways involved in protein translation, glucose, amino acid, and lipid metabolism, and HIF-1 in M1 macrophages. These results demonstrate that A3A enhances the pro-inflammatory responses in M1 macrophages, likely at the expense of genes mediating metabolic responses. Since A3A is located in the cytoplasm without translocating to the nucleus in normal monocytic cells (Land et al., 2013), and no evidence yet exists that A3A can selectively deaminate nuclear DNA in normal cells, these findings strongly suggest that its RNA editing function is responsible metabolic, gene expression, cytokine secretion and surface marker expression changes associated with M1 polarization. Since A3A-mediated RNA editing is also induced by cellular crowding and hypoxia in normal monocytes (Sharma et al., 2015), we hypothesize that C>U RNA editing plays a general role in monocyte/macrophage stress response.

Cell stress is commonly encountered by innate immune cells in pathologic microenvironments of infection and inflammation. Sudden environmental changes can induce cell stress, which triggers pathways for homeostasis or cell death, depending on the degree of cellular damage. Cells have adaptive stress responses including DNA repair, metabolic reprogramming, and UPR pathways that enable survival during states of emergency, such as infections and hypoxia (Senft and Ronai 2015). Constitutive activation of IFN-1 signaling can cause tissue dysfunction and organ failure in certain diseases, such as Aicardi-Goutières Syndrome (Rice et al., 2012). Our results support distinct roles for A3A in the adaptation of macrophages to specific stressors. In response to stressors LPS+IFN-γ, IFN-1 or viruses, A3A augments M1 polarization and pro-inflammatory responses (**Figures 2, 5 and 6**). Our results raise the speculation that while in some settings A3A may augment macrophage-mediated host defense through amplification of inflammation, A3A may be deleterious in other conditions associated with inflammatory injury, such as sepsis. While a shift from oxidative phosphorylation to aerobic glycolysis occurs in leukocytes during the initial host response to sepsis, a generalized metabolic defect involving both glycolysis and oxidative metabolism characterizes later stages of sepsis (Cheng et al., 2016). Seen in this light, our results point to A3A as a potential contributor to defects in energy metabolism associated with sepsis.

We also would like to draw some functional parallels between AID (activation-induced cytidine deaminase), encoded by AICDA, and APOBEC3A in immunity. AID and APOBEC3A are evolutionarily related cytidine deaminases (Conticello 2008) that physiologically alter the genomically encoded cellular DNA or RNA sequences as part of adaptive and innate immunity responses, respectively. AID is specifically expressed in germinal center B cells and mediates somatic hypermutation of Ig DNA, leading to generation of higher affinity antibodies in terminally differentiated memory B cells or plasma cells (Zan and Casali 2013). Whereas APOBEC3A is primarily expressed in monocyte/macrophages and its inducible RNA editing function promotes M1 pro-inflammatory phenotype in pathologic microenvironments.

The mechanism by which cellular RNA editing by A3A promotes M1 polarization requires further investigation. C>U RNA editing by A3A may mutate certain gene products or affect the stability or translatability of mRNAs, which in turn causes a cascade of downstream effects to facilitate the M1 phenotype polarization. Using high-coverage RNA_Seq data, we find a greater number of exonic/UTR C>U RNA editing sites (n=209) than initially suggested by our analysis of publicly available RNA_Seq data (n=120) in M1 macrophages (Sharma et al., 2015), including the novel identification of *THOC5* as the most (deleteriously) edited gene. *THOC5* encodes a member of the THO complex that is part of mRNA transcription/export complex (Katahira et al., 2009).

THOC5 protein is phosphorylated by several tyrosine kinases such as the M-CSF receptor (Carney et al., 2009). Tran et al. (Tran et al., 2013) showed that depletion of *THOC5* in mouse bone marrow-derived macrophages downregulated 99 genes and suppressed M-CSF mediated M2-like macrophage differentiation by inhibiting the export of M-CSF inducible genes. It is therefore conceivable that one mechanism by which RNA editing A3A promotes M1 is through physiologic suppression of THOC5, which leads to reduced M-CSF regulated gene export. Our study opens new avenues of investigation of the molecular basis by which A3A regulates inflammation and metabolism in macrophages and the potential of A3A as a therapeutic target for diseases associated with pathologic inflammation.

## Materials and Methods

### Macrophage generation and polarization

Normal human donor CD14+ monocytes were isolated from TRIMA leukoreduction filters following platelet apheresis at the blood donor center at Roswell Park Comprehensive Cancer Center (RPCCC). Since these cells were remnant products of platelet apheresis and contained no identifiers, their use was considered non-human research. Mononuclear cells were recovered with lymphocyte separation media and sepMate tubes (Stemcell Technology, Vancouver, Canada) by centrifugation, and monocytes were isolated by column purification using CD14 microbeads (Miltenyi Biotec, Somerville, MA). Cells were cultured for 1 week at a density of 10^6^/well in 6 well plates with 50 ng/ml recombinant human macrophage colony stimulating factor (M-CSF) (Life Technologies, Carlsbad, CA), 1x GlutaMAX-I (Life Technologies) and 1mM sodium pyruvate (Mediatech, Manassas, VA) to generate M0 macrophages. For M1 or M2 macrophage polarization, M0 cells were treated for two days with 20 ng/ml recombinant human IFN-y (Life Technologies) and 50 ng/ml Escherichia coli LPS (SigmaAldrich, St. Louis, MO), or 20 ng/ml recombinant human IL4 (Life Technologies), respectively. RNA was isolated from cells using TRIzol™ Reagent (Invitrogen, Carlsbad, CA) (Sharma et al., 2015).

### Knockdown of A3A in macrophages

A day before induction of M1 polarization, M0 macrophages were transfected with 100 nM of negative control (Silencer negative control no. 1, product number AM4611, Life Technologies) or equimolar mix of two human APOBEC3A siRNAs (Silencer 45715 and 45810, respectively, with sense sequences 5’-GACCUACCUGUGCUACGAATT-3’ and 5’-GCAGUAUGCUCCCGAUCAATT-3’, Life Technologies) using Lipofectamine RNAiMAX (Life Technologies), following the manufacturer’s protocol. IFN-γ and LPS were added with 2ml medium to each well, to induce M1 polarization, and cells were harvested at 48h (Sharma et al., 2015).

### Reverse transcription and PCR

RNA was reverse-transcribed with random DNA hexamers and oligo-dT primers using material and methods provided with the Transcriptor First Strand cDNA Synthesis (Roche, Indianapolis, IN). PCR typically employed 35 cycles of amplification and an annealing temperature of 60°C. Primers (Integrated DNA Technologies Coralville, IA) used for PCR of cDNA templates were designed such that the amplicons spanned multiple exons. A blend of Taq and high-fidelity Deep VentR DNA polymerases (OneTaq, New England Biolabs) was used in PCR to generate products for Sanger sequencing. Knockdown of A3A is confirmed by RT-qPCR using allele-specific primers and *SDHB* probe on a Light Cycler 480 System (Roche), as previously described (Baysal et al. 2013 and Sharma et al. 2015). Quantification cycle (Cq) values were calculated by the instrument software using the maximum second derivative method, and the mean Cq value of duplicate or triplicate PCR reactions was used for analysis (Sharma et al., 2015).

### Viral infections

#### Human monocyte-derived macrophage influenza exposure

Monocytes were isolated from freshly drawn blood, suspended in 1x GlutaMAX-I (Life Technologies) and 1mM sodium pyruvate (Mediatech) medium at 3.3×10^5^ cells/ml, and 3 ml was dispensed into each well of 6-well tissue culture plates. M-CSF, 50 ng/ml (Life Technologies), was added and the cultures were incubated at 37°C, 95% relative humidity, and 5% CO2 for 7 days. Influenza inoculum was prepared by thawing a stock vial of influenza A/Brisbane/10/2007 (H3N2) (a generous gift from Dr. Suryaprakash Sambhara at the Center for Disease Control and Prevention, Atlanta, GA), sonicated for 45 seconds in an ultrasonic water bath (Branson 1210, VWR, Radnor, PA), and diluted to 2×10^5^ PFU/ml (as determined in MDCK plaque assay) with ice cold PBS. The plated cells were washed 2X with 1 ml warm 3% BSA in DMEM after which 0.3 ml was added to prevent drying of the cells. 100 μl of influenza inoculum (MOI=0.02) or PBS control was added to appropriate wells and adsorbed for 1 hr at 37°C, 95% relative humidity, and 5% CO_2_ while gently rocking. Warm medium, 2.6 ml, was added to each well and incubation continued. After 24 or 48 hr, the medium was collected, centrifuged at 500 x g for 5 min at 4°C, and the supernatant was quick frozen in ethanol and dry ice and stored at −80°C for influenza MDCK plaque assay analysis. The cells remaining in the well were harvested by adding one ml of Trizol to each well, incubating at room temperature for 5 min and transferring to a microfuge tube and storing at −80°C for RNA analysis.

#### Influenza MDCK plaque assay

MDCK cells (ATTC, Manassas, VA) were seeded on 6-well tissue culture plates at 4×10^5^ cells/well in 2 ml of MEM with Earle’s salts + 0.1 mM non-essential amino acids + 1 mM pyruvate + 50 U/ml penicillin + 50 μg/ml + streptomycin + 20 μg/ml gentamicin+10% fetal calf serum and incubated at 37°C, 95% relative humidity, and 5% CO_2_. When cells reached 80-90% confluency (≈3 days) the cells were rinsed 2X with 1 ml 0.3% BSA in DMEM and 0.3 ml was added to prevent drying of the cells. Influenza samples were prepared, as described above, 10-fold serially diluted in 0.3% BSA in DMEM and kept on ice. Diluted samples were added to appropriate wells, 100 μl/well, and adsorbed for 1 hr at 37°C, 95% relative humidity, and 5% CO_2_ while gently rocking. The samples were aspirated and the cells rinsed 1X with PBS + 50 U/ml penicillin + 50 μg/ml + streptomycin. 2X L-15 (SigmaAldrich) with 25 mM HEPES + 0.15% NaHCO_3_ + 100 U/ml penicillin + 100 μg/ml + streptomycin + 0.5 μg/ml gentamicin + 2 μg/ml TPCK-trypsin (SigmaAldrich) was combined with an equal volume of liquified 1.0% agarose in water then 2 ml/well was dispensed into the culture wells. After incubating the plates for 2 days at 37°C, 95% relative humidity, and 5% CO_2_ the L-15/agar was removed and the cells fixed with 2 ml/well 90% ethanol for 30 min on an orbital shaker at room temperature. The ethanol was replaced with 2 ml/well 0.3% crystal violet (SigmaAldrich) in 5% isopropanol +5% ethanol in water and incubated for 20 min at room temperature on an orbital shaker. Finally, the crystal violet was removed, the cells rinsed with 2 ml/well of water, and the cultures allowed to air dry prior to counting the virus plaques.

#### Human monocyte-derived macrophage Maraba exposure

The attenuated strain MG1 Maraba virus has been previously described (Brun et al., 2010; Pol et al., 2014). Maraba virus was prepared and titered at McMaster University, shipped on dry ice to RPCCC, stored at −□80□°C and used as described (McGray et al., 2019).

### ELISA

Cytokines production were measured in supernatants of the cultured cells using ELISA kits for TNF-α, IL-1ß, IFN-α and IL6 (R&D Systems, Minneapolis, MN) following the instructions of the manufacturer.

### Flow cytometry

The following antibodies were used: Anti-human CD33 PE-Cyanine7 (eBioscience, San Diego, CA), anti-human CD86 APC (BioLegend, San Diego, CA), and antihuman CD206 FITC (BioLegend). Annexin V FITC (BioLegend) and propidium iodide solution (Sigma) were used for detection of cell death and apoptosis. Flow cytometry analysis was conducted on a LSRII flow cytometer (Becton Dickinson, Franklin Lakes, NJ). Forward scatter versus side scatter gating was set to include all non-aggregated cells from at least 20,000 events collected per sample and data were analyzed using Winlist 3D version 8.0 (Verity, Topsham ME).

### Seahorse assay

For all extracellular flux assays, cell were plated on cell-take coated Seahorse XF96 cell culture microplates at a density of 0.5 X10^5^ cells per well. The assay plates were spun for 5 minutes at 1,000 rpm and incubated at 37°C without CO_2_ before performing the assay on the Seahorse Bioscience XFe96. The Mitochondrial Stress Test was conducted in XF media containing 10mM glucose, 1mM sodium pyruvate, and 2mM L-glutamine and the following inhibitors were added at the final concentrations: Oligomycin (2μM), Carbonyl cyanide 4-(trifluoromethoxy) phenylhydrazone (FCCP) (2 μM), Rotenone/Antimycin A (0.5μM each). The Glycolytic Stress Test was performed in XF media containing 2mM L-glutamine, and the following reagents were added at the final concentrations: Glucose (10mM), Oligomycin (2μM), and 2-deoxyglucose (50mM). The data was analyzed using Wave software.

### Tumor-associated macrophage (TAM) isolation

Ascites was collected at the time of primary surgery from patients with newly diagnosed stage III epithelial ovarian cancer (EOC) under an IRB-approved protocol. All subjects signed informed consent before surgery. Ascites cells were pelleted by centrifugation (500g for 10 min at 4°C). Cells were either used within 24h of harvest for flow cytometry and functional studies or frozen in liquid nitrogen in media containing 20% FBS and 5% DMSO. Ascites macrophages were purified by CD14 microbeads, and purity was confirmed by flow cytometry demonstrating CD33+^i^ cells and by cytology (Khan et al., 2015). Universal type IFN-1, a hybrid of amino-terminal IFNα-2 and carboxy-terminal IFNα-1, produced in *E*. *Coli,* is used to evaluate RNA editing induction in TAMs and was obtained from PBL Assay Science (Piscataway, NJ).

### RNA sequencing for A3A scramble (SC) vs A3A knockdown (KD) transfected macrophages

#### Differential expression analysis of RNA data

We used a similar approach for RNA_Seq analysis as previously described (Sharma et al., 2019). Three experimental conditions including M0 macrophages, control scrambled siRNA (SC) and A3A siRNA (KD) are used. SC and KD samples were derived from the same 3 donors, whereas the M0 macrophages were derived from 3 unrelated donors. RNA libraries were prepared using the Illumina TruSeq RNA Exome protocol. For each sample, a total of ~90 million paired-end reads were obtained and QCed using fastqc (v0.10.1) (Andrews 2010). The reads were mapped to GRCh38 human reference genome and GENCODE (v25) annotation database using TopHat (v2.0.13) (Trapnell et al., 2009) with a maximum of 1 mismatch per read. The aligned reads were further checked with RSeQC (v2.6.3) (Wang et al., 2012) in order to identify potential RNASeq related problems. Gene level read counts were estimated with featureCounts from Subread (v1.6.0) (Liao et al., 2014) using --fracOverlap 1 option. Differential expression analyses were performed using DESeq2 (v1.18.1) (Love et al., 2014) and the heatmaps were generated using pheatmap (v1.0.8) R library (Kolde and Kolde 2015). Pathways analyses were performed running GSEA (v3.0b3) (Subramanian et al., 2005) pre-ranked mode with genes ordered by DESeq2’s test statistic and MiSigDB canonical curated gene sets (c2).

#### RNA editing detection

RNA Reads were mapped to reference genome GRCh38 using a lenient alignment strategy allowing at most 2 mismatches per read. Each mapped sample was run through the GATK best practices pipeline to obtain a list of variant sites. A filtered list of candidate editing events was collected based on selecting only variants that match the editing event of interest (i.e. C>T along with G>A and A>G along with T>C). Returning to the mapping results, samples are ‘piledup’ on the union of all potential candidates. Piledup samples were formatted and compiled for statistical comparisons based on a series of specific conditional filtering. Potential candidates for RNA editing required the following features: A) variant spots with at least two piledup records per group (SC, KD or M0) and B) at least two samples from the experimental condition SC have > 5% of editing level (el). A generalized linear model (GLM) was used to model and call editing events within the populations. The number of alternative nucleotides was compared to the reference between the two groups (Group). The first filter was designed to remove any possible known SNPs. The resulting editing candidates were annotated according to dbSNP144. A first layer of filtering removed those events where at least one sample had > 0.95 event level, corresponding to homozygous SNPs. Then a second layer was performed where an event was removed if all of the samples had any of the following SNP features: el<0.05; 0.4<el<0.6 and el>0.95 and if this event was matched in dbSNP database. The next filtering step removed all C>T (or G>A) events that were not preceded by C or T in the + strand and G or A in the – strand. A resulting table specifying the editing site, the type of editing event, editing level and number of reads on a reference and alternative bases on each sample for each group was generated. The editing events were annotated using ANNOVAR’s gene-based annotation to identify gene features and protein amino acid sequence changes. Finally, a manual stringency filter was employed to retain only the exonic or UTR sites where (a) the editing level increases 2-fold or more in SC relative to M0 and (b) the edited C is located at the 3’-end of a putative tri- or tetra loop that is flanked by a stem that has at least 2 bp long perfect complementarity, or at least 4 bp long imperfect complementarity in which 1 nucleotide mismatch or bulging was allowed (Sharma and Baysal 2017).

### Sanger sequencing

PCR-amplified cDNA fragments were sequenced to examine candidate RNA-editing sites. PCR reactions were treated with an ExoSAP-IT exonuclease (Affymetrix, Santa Clara, CA) and then directly used for sequencing on 3130 XL Genetic Analyzer (Life Technologies) via RPCCC Genomic Core Facility. Major and minor chromatogram peak heights at a nucleotide position of interest were visualized using Sequencher 5.0 software (Gene Codes, Ann Arbor, MI).

### Sequence deposition in a public repository

Sequence data from RNA_Seq experiments are deposited to GEO repository at https://www.ncbi.nlm.nih.gov/geo/ under accession number GSE146867.

## Acknowledgements

This research was supported by educational funds from Saudi Arabia (EAQ), startup funds from the Departments of Pathology (BEB), and NIH grant R01CA188900 (BHS) and by National Cancer Institute (NCI) Grant (P30CA016056) involving the use of RPCCC’s Genomics Shared Resources, Bioinformatics Shared Resources, Flow Cytometry and Imaging, Immune Analysis Facilities and Clinical Research Services. JW and ECG are also supported by U24CA232979.

## Author Contributions

BEB and BHS conceived the study and designed the experiments with contributions from EYA. EYA performed most of the experiments with contributions from SS (RNA editing), ANHK (cell purification and culture), TE (cell purification and culture), KLS (cell purification and culture), BAD (Influenza virus growth and maintenance), QL (statistical support) and JM, AJRM and BDL (Maraba virus growth and maintenance) with support from KO. KBM and KO contributed to experimental planning. EYA, BHS and BEB wrote the manuscript. EYA and BEB prepared the figures. ECG and JW analyzed the RNA-seq data and wrote the method. All authors read and approved the final manuscript.

---

**Supp. Figure 1.**
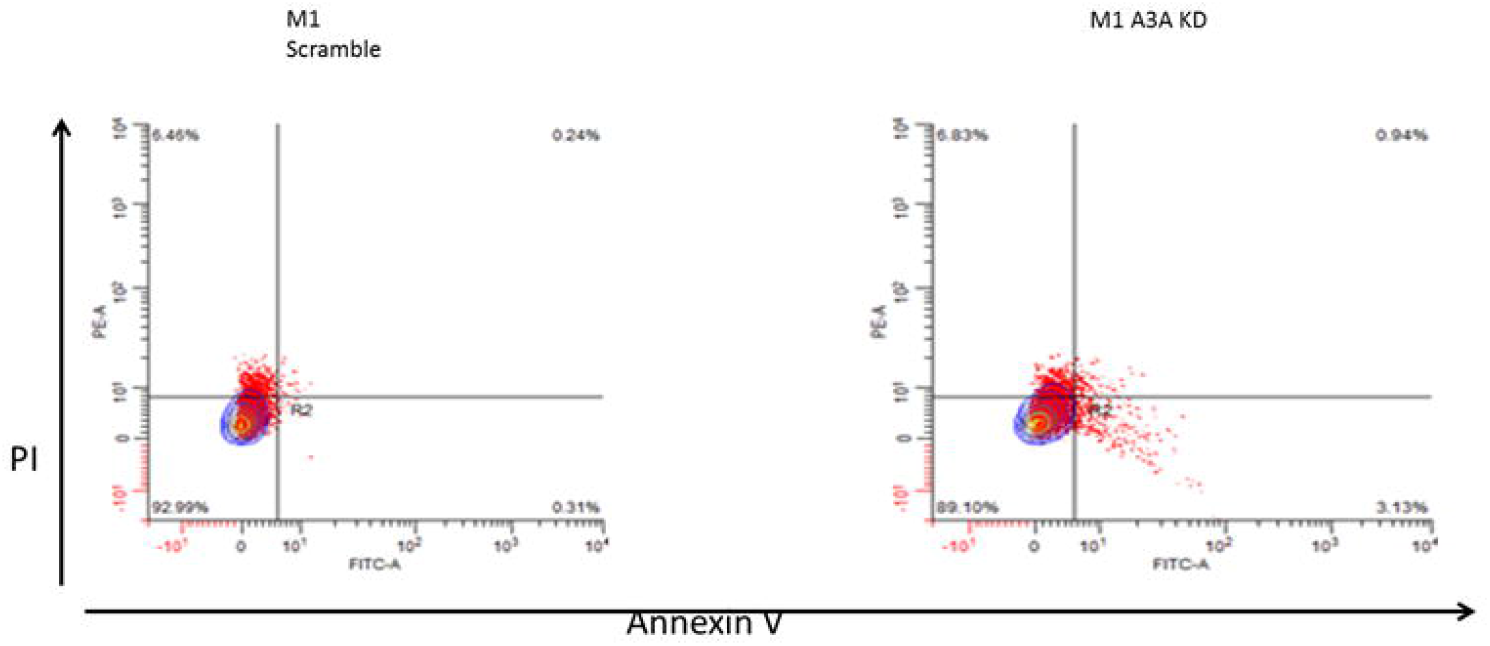
A3A does not affect cell viability in Ml-macrophages. Monocytes isolated from normal donor blood were differentiated to MO macrophages as described. MO macrophages, after A3A knock down (or control siRNA transfection, SC), were polarized to Ml macrophages with IFN-Ï (20ng/ml for 48h) and LPS (50ng/ml for 48h) and evaluated by flow cytometry for cell death using Annexin V and PI. Percent viable cells are similar in KD (93%) and SC (89%).

## Notes

### Competing Interest Statement

The authors have declared no competing interest.

### Summary of Updates

(1) Revised statistical analysis and Figure 6E (2) addition of co-author (Q.Liu) (3) other minor changes in text

## References

Andrews S. 2010. FastQC: a quality control tool for high throughput sequence data. Babraham Bioinformatics, Babraham Institute, Cambridge, United Kingdom

Baysal BE. 2007. A recurrent stop-codon mutation in succinate dehydrogenase subunit B gene in normal peripheral blood and childhood T-cell acute leukemia. PLoS One 2: e436 doi:10.1371/journal.pone.0000436.

Baysal BE, De Jong K, Liu B, Wang J, Patnaik SK, Wallace PK, Taggart RT. 2013. Hypoxia-inducible C-to-U coding RNA editing downregulates SDHB in monocytes. PeerJ 1: e152 doi:10.7717/peerj.152;10.7717/peerj.152.

Bransteitter R, Prochnow C, Chen XS. 2009. The current structural and functional understanding of APOBEC deaminases. Cell Mol Life Sci 66: 3137–3147 doi:10.1007/s00018-009-0070-y.

Breitbach CJ. 2020. Considerations for Clinical Translation of MG1 Maraba Virus. Methods in molecular biology (Clifton, NJ) 2058: 285–293 doi:10.1007/978-1-4939-9794-7_19.

Brun J, McManus D, Lefebvre C, Hu K, Falls T, Atkins H, Bell JC, McCart JA, Mahoney D, Stojdl DF. 2010. Identification of genetically modified Maraba virus as an oncolytic rhabdovirus. Molecular Therapy 18: 1440–1449 doi:10.1038/mt.2010.103.

Carney L, Pierce A, Rijnen M, Gonzalez Sanchez MB, Hamzah HG, Zhang L, Tamura T, Whetton AD. 2009. THOC5 couples M-CSF receptor signaling to transcription factor expression. Cell Signal 21: 309–316 doi:10.1016/j.cellsig.2008.10.018.

Cheng SC, Scicluna BP, Arts RJ, Gresnigt MS, Lachmandas E, Giamarellos-Bourboulis EJ, Kox M, Manjeri GR, Wagenaars JA, Cremer OL, Leentjens J, van der Meer AJ, van de Veerdonk FL, Bonten MJ, Schultz MJ, Willems PH, Pickkers P, Joosten LA, van der Poll T, Netea MG. 2016. Broad defects in the energy metabolism of leukocytes underlie immunoparalysis in sepsis. Nat Immunol 17: 406–413 doi:10.1038/ni.3398.

Conticello SG. 2008. The AID/APOBEC family of nucleic acid mutators. Genome Biol 9: 229 doi:10.1186/gb-2008-9-6-229.

Edwards JP, Zhang X, Frauwirth KA, Mosser DM. 2006. Biochemical and functional characterization of three activated macrophage populations. J Leukoc Biol 80: 1298–1307 doi:10.1189/jlb.0406249.

Eisenberg E, Levanon EY. 2018. A-to-I RNA editing—immune protector and transcriptome diversifier. Nature Reviews Genetics 19: 473 doi:10.1038/s41576-018-0006-1.

Farajollahi S, Maas S. 2010. Molecular diversity through RNA editing: a balancing act. Trends in Genetics 26: 221–230 doi:10.1016/j.tig.2010.02.001.

Galván-Peña S, O’Neill LA. 2014. Metabolic reprograming in macrophage polarization. Frontiers in immunology 5: 420 doi:10.3389/fimmu.2014.00420.

Grünewald J, Zhou R, Iyer S, Lareau CA, Garcia SP, Aryee MJ, Joung JK. 2019. CRISPR DNA base editors with reduced RNA off-target and self-editing activities. Nature biotechnology 37: 1041–1048 doi:10.1038/s41587-019-0236-6.

Holmes RK, Malim MH, Bishop KN. 2007. APOBEC-mediated viral restriction: not simply editing? Trends in biochemical sciences 32: 118–128 doi:10.1016/j.tibs.2007.01.004.

Katahira J, Inoue H, Hurt E, Yoneda Y. 2009. Adaptor Aly and co-adaptor Thoc5 function in the Tap-p15-mediated nuclear export of HSP70 mRNA. EMBO J 28: 556–567 doi:10.1038/emboj.2009.5.

Khan AN, Kolomeyevskaya N, Singel KL, Grimm MJ, Moysich KB, Daudi S, Grzankowski KS, Lele S, Ylagan L, Webster GA, Abrams SI, Odunsi K, Segal BH. 2015. Targeting myeloid cells in the tumor microenvironment enhances vaccine efficacy in murine epithelial ovarian cancer. Oncotarget 6: 11310–11326 doi:10.18632/oncotarget.3597.

Kolde R, Kolde MR. 2015. Package ‘pheatmap’. R Package 1.

Krishnan V, Schaar B, Tallapragada S, Dorigo O. 2018. Tumor associated macrophages in gynecologic cancers. Gynecol Oncol 149: 205–213 doi:10.1016/j.ygyno.2018.01.014.

Land AM, Law EK, Carpenter MA, Lackey L, Brown WL, Harris RS. 2013. Endogenous APOBEC3A DNA cytosine deaminase is cytoplasmic and nongenotoxic. The Journal of biological chemistry 288: 17253–17260 doi:10.1074/jbc.M113.458661 [doi].

Laskin DL. 2009. Macrophages and inflammatory mediators in chemical toxicity: a battle of forces. Chem Res Toxicol 22: 1376–1385 doi:10.1021/tx900086v.

Lawrence T, Natoli G. 2011. Transcriptional regulation of macrophage polarization: enabling diversity with identity. Nature reviews Immunology 11: 750–761 doi:10.1038/nri3088.

Liao Y, Smyth GK, Shi W. 2014. featureCounts: an efficient general purpose program for assigning sequence reads to genomic features. Bioinformatics 30: 923–930 doi:10.1093/bioinformatics/btt656.

Love MI, Huber W, Anders S. 2014. Moderated estimation of fold change and dispersion for RNA-seq data with DESeq2. Genome Biol 15: 550 doi:10.1186/s13059-014-0550-8.

Martin NT, Bell JC. 2018. Oncolytic Virus Combination Therapy: Killing One Bird with Two Stones. Molecular therapy: the journal of the American Society of Gene Therapy 26: 1414–1422 doi:10.1016/j.ymthe.2018.04.001.

McGray AJR, Huang R-Y, Battaglia S, Eppolito C, Miliotto A, Stephenson KB, Lugade AA, Webster G, Lichty BD, Seshadri M, Kozbor D, Odunsi K. 2019. Oncolytic Maraba virus armed with tumor antigen boosts vaccine priming and reveals diverse therapeutic response patterns when combined with checkpoint blockade in ovarian cancer. Journal for immunotherapy of cancer 7: 189 doi:10.1186/s40425-019-0641-x.

Mills EL, Kelly B, Logan A, Costa ASH, Varma M, Bryant CE, Tourlomousis P, Dabritz JHM, Gottlieb E, Latorre I, Corr SC, McManus G, Ryan D, Jacobs HT, Szibor M, Xavier RJ, Braun T, Frezza C, Murphy MP, O’Neill LA. 2016. Succinate Dehydrogenase Supports Metabolic Repurposing of Mitochondria to Drive Inflammatory Macrophages. Cell 167: 457–470 e413 doi:10.1016/j.cell.2016.08.064.

O’Neill LA, Pearce EJ. 2016. Immunometabolism governs dendritic cell and macrophage function. Journal of Experimental Medicine 213: 15–23 doi:10.1084/jem.20151570.

Pol JG, Zhang L, Bridle BW, Stephenson KB, Rességuier J, Hanson S, Chen L, Kazdhan N, Bramson JL, Stojdl DF. 2014. Maraba virus as a potent oncolytic vaccine vector. Molecular Therapy 22: 420–429 doi:10.1038/mt.2013.249.

Porcheray F, Viaud S, Rimaniol AC, Leone C, Samah B, Dereuddre-Bosquet N, Dormont D, Gras G. 2005. Macrophage activation switching: an asset for the resolution of inflammation. Clinical and experimental immunology 142: 481–489 doi:10.1111/j.1365-2249.2005.02934.x.

Rice GI, Kasher PR, Forte GM, Mannion NM, Greenwood SM, Szynkiewicz M, Dickerson JE, Bhaskar SS, Zampini M, Briggs TA, Jenkinson EM, Bacino CA, Battini R, Bertini E, Brogan PA, Brueton LA, Carpanelli M, De Laet C, de Lonlay P, del Toro M, Desguerre I, Fazzi E, Garcia-Cazorla A, Heiberg A, Kawaguchi M, Kumar R, Lin JP, Lourenco CM, Male AM, Marques W, Jr., Mignot C, Olivieri I, Orcesi S, Prabhakar P, Rasmussen M, Robinson RA, Rozenberg F, Schmidt JL, Steindl K, Tan TY, van der Merwe WG, Vanderver A, Vassallo G, Wakeling EL, Wassmer E, Whittaker E, Livingston JH, Lebon P, Suzuki T, McLaughlin PJ, Keegan LP, O’Connell MA, Lovell SC, Crow YJ. 2012. Mutations in ADAR1 cause Aicardi-Goutieres syndrome associated with a type I interferon signature. Nat Genet 44: 1243–1248 doi:10.1038/ng.2414.

Rodríguez-Prados J-C, Través PG, Cuenca J, Rico D, Aragonés J, Martín-Sanz P, Cascante M, Boscá L. 2010. Substrate fate in activated macrophages: a comparison between innate, classic, and alternative activation. The Journal of Immunology 185: 605–614 doi:10.4049/jimmunol.0901698.

Röszer T. 2015. Understanding the mysterious M2 macrophage through activation markers and effector mechanisms. Mediators of inflammation 2015 doi:10.1155/2015/816460.

Salter JD, Bennett RP, Smith HC. 2016. The APOBEC Protein Family: United by Structure, Divergent in Function. Trends Biochem Sci 41: 578–594 doi:10.1016/j.tibs.2016.05.001.

Sang Y, Miller LC, Blecha F. 2015. Macrophage Polarization in Virus-Host Interactions. J Clin Cell Immunol 6 doi:10.4172/2155-9899.1000311.

Senft D, Ronai ZA. 2015. UPR, autophagy, and mitochondria crosstalk underlies the ER stress response. Trends Biochem Sci 40: 141–148 doi:10.1016/j.tibs.2015.01.002.

Sharma S, Baysal BE. 2017. Stem-loop structure preference for site-specific RNA editing by APOBEC3A and APOBEC3G. PeerJ 5: e4136 doi:10.7717/peerj.4136.

Sharma S, Patnaik SK, Kemer Z, Baysal BE. 2017a. Transient overexpression of exogenous APOBEC3A causes C-to-U RNA editing of thousands of genes. RNA Biol 14: 603–610 doi:10.1080/15476286.2016.1184387.

Sharma S, Patnaik SK, Taggart RT, Baysal BE. 2016. The double-domain cytidine deaminase APOBEC3G is a cellular site-specific RNA editing enzyme. Scientific reports 6: 39100 doi:10.1038/srep39100 [doi].

Sharma S, Patnaik SK, Taggart RT, Kannisto ED, Enriquez SM, Gollnick P, Baysal BE. 2015. APOBEC3A cytidine deaminase induces RNA editing in monocytes and macrophages. Nat Commun 6: 6881 doi:10.1038/ncomms7881.

Sharma S, Wang J, Alqassim E, Portwood S, Cortes Gomez E, Maguire O, Basse PH, Wang ES, Segal BH, Baysal BE. 2019. Mitochondrial hypoxic stress induces widespread RNA editing by APOBEC3G in natural killer cells. Genome Biol 20: 37 doi:10.1186/s13059-019-1651-1.

Sharma S, Wang J, Cortes Gomez E, Taggart RT, Baysal BE. 2017b. Mitochondrial complex II regulates a distinct oxygen sensing mechanism in monocytes. Hum Mol Genet 26: 1328–1339 doi:10.1093/hmg/ddx041.

Subramanian A, Tamayo P, Mootha VK, Mukherjee S, Ebert BL, Gillette MA, Paulovich A, Pomeroy SL, Golub TR, Lander ES, Mesirov JP. 2005. Gene set enrichment analysis: a knowledge-based approach for interpreting genome-wide expression profiles. Proc Natl Acad Sci U S A 102: 15545–15550 doi:10.1073/pnas.0506580102.

Teng B, Burant CF, Davidson NO. 1993. Molecular cloning of an apolipoprotein B messenger RNA editing protein. Science 260: 1816–1819.

Tong JG, Valdes YR, Sivapragasam M, Barrett JW, Bell JC, Stojdl D, DiMattia GE, Shepherd TG. 2017. Spatial and temporal epithelial ovarian cancer cell heterogeneity impacts Maraba virus oncolytic potential. BMC Cancer 17: 594 doi:10.1186/s12885-017-3600-2.

Tran DD, Saran S, Dittrich-Breiholz O, Williamson AJ, Klebba-Farber S, Koch A, Kracht M, Whetton AD, Tamura T. 2013. Transcriptional regulation of immediate-early gene response by THOC5, a member of mRNA export complex, contributes to the M-CSF-induced macrophage differentiation. Cell Death Dis 4: e879 doi:10.1038/cddis.2013.409.

Trapnell C, Pachter L, Salzberg SL. 2009. TopHat: discovering splice junctions with RNA-Seq. Bioinformatics (Oxford, England) 25: 1105–1111 doi:10.1093/bioinformatics/btp120 [doi].

Van Ginderachter JA, Movahedi K, Ghassabeh GH, Meerschaut S, Beschin A, Raes G, De Baetselier P. 2006. Classical and alternative activation of mononuclear phagocytes: picking the best of both worlds for tumor promotion. Immunobiology 211: 487–501 doi:10.1016/j.imbio.2006.06.002.

Vats D, Mukundan L, Odegaard JI, Zhang L, Smith KL, Morel CR, Greaves DR, Murray PJ, Chawla A. 2006. Oxidative metabolism and PGC-1ß attenuate macrophage-mediated inflammation. Cell metabolism 4: 13–24 doi:10.1016/j.cmet.2006.05.011.

Wang L, Wang S, Li W. 2012. RSeQC: quality control of RNA-seq experiments. Bioinformatics 28: 2184–2185 doi:10.1093/bioinformatics/bts356.

Wolf MT, Dearth CL, Ranallo CA, LoPresti ST, Carey LE, Daly KA, Brown BN, Badylak SF. 2014. Macrophage polarization in response to ECM coated polypropylene mesh. Biomaterials 35: 6838–6849 doi:10.1016/j.biomaterials.2014.04.115.

Wynn TA, Chawla A, Pollard JW. 2013. Macrophage biology in development, homeostasis and disease. Nature 496: 445–455 doi:10.1038/nature12034.

Zan H, Casali P. 2013. Regulation of Aicda expression and AID activity. Autoimmunity 46: 83–101 doi:10.3109/08916934.2012.749244.

Zhang X, Mosser DM. 2008. Macrophage activation by endogenous danger signals. J Pathol 214: 161–178 doi:10.1002/path.2284.

Zhou D, Yang K, Chen L, Zhang W, Xu Z, Zuo J, Jiang H, Luan J. 2017. Promising landscape for regulating macrophage polarization: epigenetic viewpoint. Oncotarget 8: 57693–57706 doi:10.18632/oncotarget.17027.

